# Oct4 regulates embryonic pluripotency via metabolic mechanisms and Stat3 signalling

**DOI:** 10.1101/2020.01.28.922856

**Authors:** Giuliano Giuseppe Stirparo, Agata Kurowski, Stanley Eugene Strawbridge, Hannah Stuart, Thorsten Edwin Boroviak, Ayaka Yanagida, Jennifer Nichols

**Author notes:** These authors contributed equally (Giuliano Giuseppe Stirparo, Agata Kurowski).

## Abstract

OCT4 is a fundamental component of the molecular circuitry governing pluripotency *in vivo* and *in vitro*. To determine how OCT4 protects the pluripotent lineage from differentiation into trophoblast, we used single cell transcriptomics and quantitative immunofluorescence on blastocysts and established differentially expressed genes and pathways between control and OCT4 null cells. Activation of most pluripotency-associated transcription factors in the early mouse inner cell mass appears independent of OCT4, whereas JAK/STAT signalling requires OCT4, via activation of IL6ST. Single cell deconvolution, diffusion component and trajectory inference dissected the process of differentiation of OCT4 null cells by activating specific gene-network and transcription factors. Downregulation of glycolytic and oxidative metabolism was observed. CHIPseq analysis suggests OCT4 directly targets rate-limiting glycolytic enzymes. Concomitant with significant disruption of the STAT3 pathway, oxidative respiration is significantly diminished in OCT4 null cells. Upregulation of the lysosomal pathway detected in OCT4 null embryos is likely attributable to aberrant metabolism.

**Highlights and novelty:**

- Major pluripotency-associated transcription factors are activated in OCT4-deficient early mouse ICM cells, coincident with ectopic expression of trophectoderm markers
- JAK/STAT signalling is defective in OCT4 null embryos
- OCT4 promotes expression of KATS enzymes by means of glycolytic production of Acetyl CoA to secure chromatin accessibility for acquisition of epiblast identity
- OCT4 regulates the metabolic and biophysical processes required for establishment of embryonic pluripotency

## Introduction

Formation of a mammalian organism pivots upon establishment of extraembryonic tissues to pattern the foetus and expedite connection with the maternal vascular system, whilst preserving a pluripotent population of cells with the responsive capacity to generate body pattern and tissues progressively during development. Specification of trophectoderm (TE, founder of the placenta) on the outside of the preimplantation embryo coincides with appearance of the blastocyst cavity and a metabolic switch from pyruvate and lactose to glucose utilisation with increased oxygen consumption^1–5^. This dramatic change heralds an increase in metabolic activity by the differentiating TE, comprising elevated ATP, amino acid turnover and mitochondrial count^6, 7^. The murine embryo is equipped to overcome adverse consequences associated with accumulation of reactive oxygen species during the metabolic transition to oxidative phosphorylation, largely facilitated by the transcriptional enhancer factor TEAD4^8, 9^. TEAD4 becomes intensified in the TE, where it cooperates with nuclear YAP to initiate transcription of TE-specific genes^10, 11^. TE differentiation is orchestrated by a specific set of transcription factors; acquisition of TE identity actuates distinct metabolic requirements compared with the undifferentiated inner cell mass (ICM). During blastocyst expansion, the transcription factor OCT4 (encoded by *Pou5f1*) becomes restricted to the ICM^12^. OCT4 is essential for establishment of the pluripotent epiblast, preventing differentiation of the embryo towards TE^13^, and is absolutely required for propagation of pluripotent stem cells *in vitro*^13–17^. Studies in embryonic stem cells (ESC) indicate that the pluripotency network hinges upon OCT4^18–22^. In the embryo OCT4 is detected throughout cleavage^12^, whereas many other pluripotency-associated factors, such as NANOG, appear after the onset of zygotic genome activation^23^. However, in embryos lacking both maternal and zygotic OCT4, NANOG emerges robustly, coincidentally with OCT4-positive controls^24, 25^, ruling out failure to express this key pluripotency network gene as a contributing feature of the OCT4 null phenotype. Whether absence of other pluripotency-associated factors could play a role has not been investigated at the single cell level. To date, evidence that the cells occupying an inside position in OCT4 null embryos adopt a TE identity is largely restricted to morphology and expression of TE-specific markers at the time of implantation^13, 24, 26^. To scrutinise the apparent lineage transition and probe the mechanism by which acquisition of pluripotency in OCT4 null ICMs fails we used single cell RNA sequencing (scRNAseq) and quantitative immunofluorescence (QIF) to examine gene expression in wild type, heterozygous and OCT4 null early and late blastocyst ICMs. Differences between samples and groups, calculated using bioinformatics and computational analysis, revealed novel insight into the role of OCT4 in defining the metabolic, pluripotent and biophysical status of the murine ICM.

## Results

### Single cell transcriptional profiling reveals divergence of OCT4 null from wild type and heterozygous ICM cells by the mid blastocyst stage

Following observation of NANOG protein in OCT4 null blastocysts^24, 25^, we performed whole transcriptome analysis by scRNAseq of ICM cells isolated from *Pou5f1* heterozygous *inter se* mating at embryonic day (E)3.5 to investigate the entire pluripotency network. Quality control, as previously reported^27^, eliminated inadequate samples, leaving 29 null mutant (MUT), 42 wild-type (WT) and 16 heterozygous (HET) cells from 4, 5 and 2 mid blastocysts, respectively (Fig.1A, Suppl.Table1). *Pou5f1* RNA expression was not detected in MUT embryos, confirming degradation of maternal transcripts (Fig.1A, Fig.S1A), consistent with lack of OCT4 protein observed at the morula stage^13^. To characterize global differences and similarities between genotypes t-SNE analysis was performed (Fig.1B, Fig.S1A) using the most variable genes identified in early blastocysts (n=2232, log_2_FPKM>0.5, logCV_2_>0.5). MUT cells cluster separately from HET and WT, suggesting major changes in transcriptome, despite no apparent difference in ICM morphology. Interestingly, HET and WT cells clustered together, indicating no more than a negligible effect of reduced *Pou5f1* in HET cells in the developing embryo, contrasting with the elevated and more homogeneous expression of *Nanog, Klf4* and Estrogen-related-receptor B *(Esrrb)* previously reported in *Oct4* HET ESCs^28^ (Fig.S1B).

**Figure 1:**
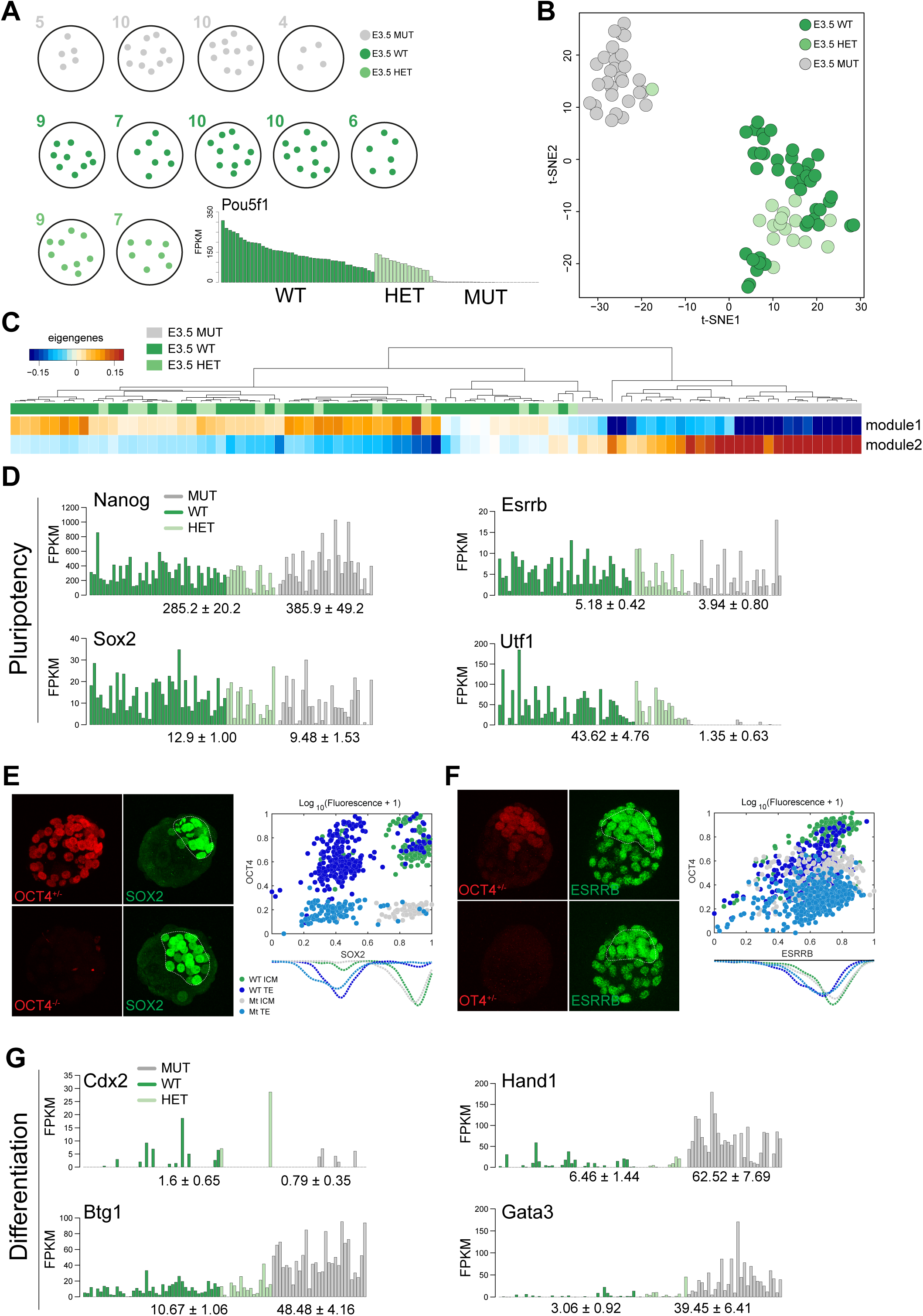
(A) Schematic of number of single cells per embryo (E3.5 stages) and their genotype. Barplot shows FPKM expression of *Pou5f1* for each single cell. (B) t-SNE plot for early blastocyst cells. Sample colour represents the different genotypes. (C) One-way hierarchical cluster of eigengenes value computed from WGCNA (power 10; dist=0.35, size =30). (D) FPKM expression of genes associated with Pluripotency. (E) Confocal images and normalized expression of OCT4 HET and MUT embryos stained for SOX2 and (F) ESRRB. (G) FPKM expression of trophectoderm markers.

Weighted gene correlation network analysis (WGCNA) allows extraction of modules defined by co-regulated genes, combined with unsupervised clustering (Fig.1C, Fig.S1C). Two main modules emerged, clustering cells according to genotype: module 1 co-clusters HET and WT and co-regulates pluripotency-associated genes such as *Pou5f1, Gdf3* and *Zfp42*^29–31^ (Fig.S1D); module 2 is specific for MUT cells, expressing established TE markers, including *Hand1, Krt18* and *Gata3* (Fig.S1D)^32–35^.

### Activation of major pluripotency network elements in the mouse blastocyst is independent of OCT4, but suppression of ectopic TE expression is not

In light of the significant transcriptional differences revealed, we sought insight into regulation of the pluripotency network in WT/HET and MUT cells (Fig.1D, Fig.S1D, Suppl.Table2). Consistent with previously published immunohistochemistry (IHC) data^24, 25^ *Nanog* was detected, albeit heterogeneously, in MUT cells (Fig.1D). Conversely, *Sox2* was not significantly affected in MUT cells at both RNA (Fig.1D) and protein levels (Fig.1E), as detected by quantitative immunofluorescence (QIF)^36^. *Esrrb*, reported to be a direct OCT4 target *in vivo*^24^, showed modest downregulation in MUT cells by scRNAseq (Fig.1D), but no obvious difference at the protein level via QIF (Fig.1F) suggesting initiation of expression independent of OCT4 in early blastocysts. Specific chromatin components are involved in establishing and maintaining pluripotency *in vivo* and *in vitro*^37^. *Utf1*, a direct target of OCT4^38^, is expressed in normal ICM and epiblast^39^; its expression decreases upon differentiation^40^, consistent with its role in maintaining chromatin structure compatible with self-renewal *in vitro*^41^. As expected, *Utf1* was not detected in MUT early blastocysts (Fig.1D). TE markers, such as *Btg1*, *Hand1* and *Gata3* were found in most MUT cells, whereas *Cdx2* was poorly represented (5/29 MUT cells; Fig.1G), suggesting that TE differentiation of MUT cells is not primarily directed by *Cdx2*, although its protein appears in the majority of OCT4 null ICMs by E4.0^26^.

### Reduction of JAK/STAT signalling machinery distinguishes OCT4 null ICMs

The JAK/STAT signalling pathway is fundamental for self-renewal and pluripotency *in vivo* and *in vitro*^42–44^. Active P-STAT3 protein and its targets *Klf4*^45^ and *Tfcp2l1*^46^ were significantly lower in MUT cells at both protein and mRNA levels (Fig.2A-D). Total *Stat3* mRNA levels did not vary (Fig. S2A). The reduced STAT3 signalling in MUT embryos was most likely attributable to absence of its upstream cytokine receptor subunit, *gp130* (*Il6st*) (Fig.2A). *Il6st* is also a putative target of OCT4 in ESC (Suppl.Table3, https://chip-atlas.org/). Unsurprisingly, Suppressor of Cytokine Signalling *(Socs)3*, a direct STAT3 target that exerts negative feedback regulation^47^ was barely detectable in MUT cells (Fig.2A). PCA computed with genes in JAK/STAT signalling pathway (https://www.genome.jp/kegg/) segregates MUT from WT/HET cells (Fig.2E); cumulative sum on the relative percentage of gene expression is significantly higher (pval < 0.05) in WT/HET, indicating that in MUT cells the pathway is downregulated (Fig.2F).

**Figure 2:**
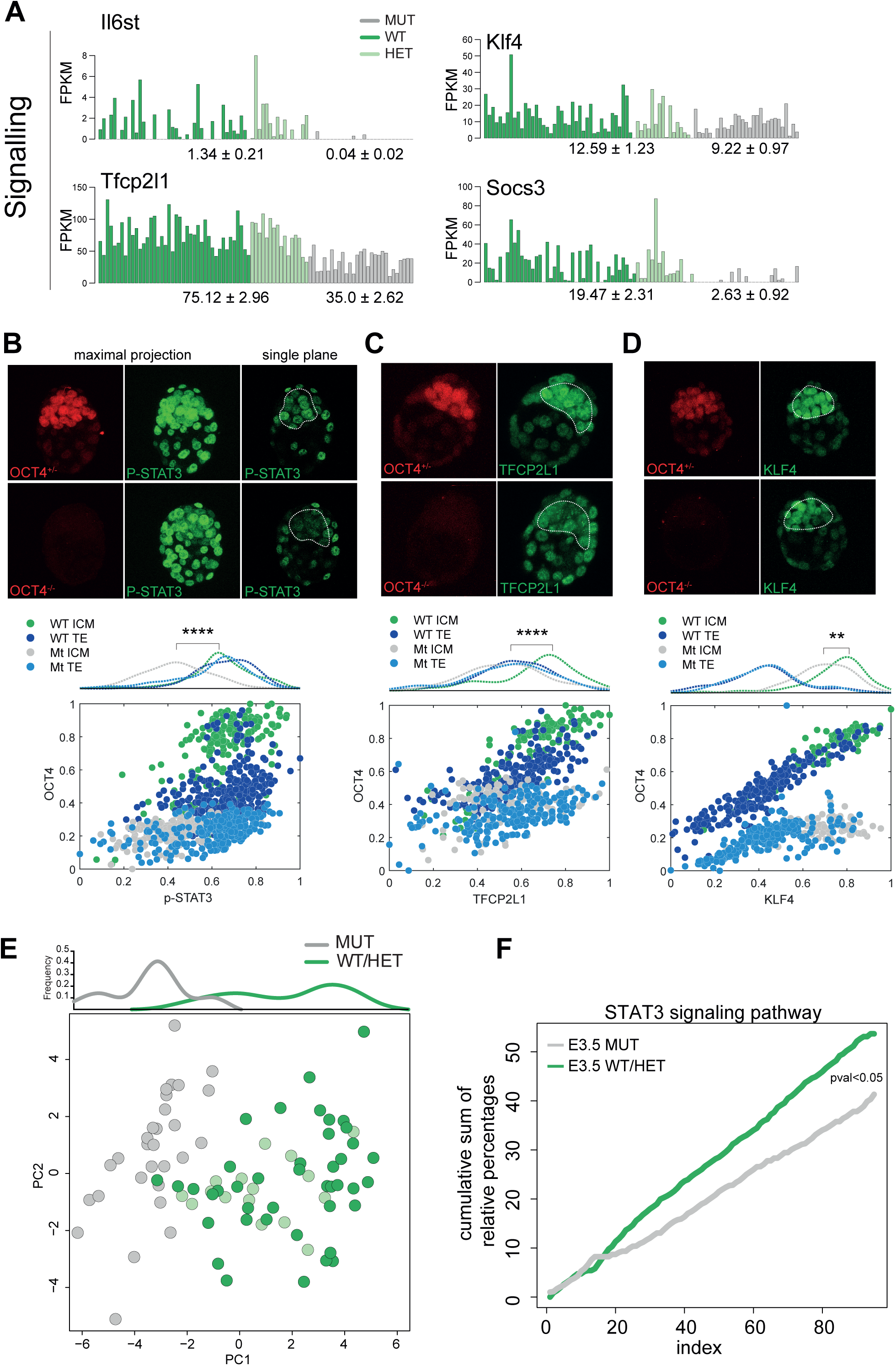
(A) FPKM expression of genes in STAT3 pathway. (B) Confocal images and normalized expression of OCT4 HET and MUT embryos stained for p-STAT3, (C) TFCP2L1, and (D) KLF4. (E) PCA computed with genes in KEGG JAK/STAT signalling pathway. (F) Cumulative sum of relative percentages between the expression of genes in STAT3 pathway (KEGG); student’s t-test p.val < 0.05.

The results presented thus far reveal reduced expression of certain factors in MUT cells, particularly direct OCT4 targets and JAK/STAT pathway members, as well as ectopic activation of some TE marker genes, indicating transcriptional divergence in MUT cells by E3.5.

### Dissecting overt impairment of lineage segregation in maturing OCT4 null ICMs

For detailed characterisation of events associated with diversion of ICM to TE in embryos lacking OCT4, diffusion component analysis was performed on implanting embryos 24 hours older (E4.5) (Fig.3A; Fig.S3A); 19 cells isolated from 2 MUT, 22 from 2 WT and 44 from 4 HET late ICMs were analysed (Suppl.Table1, Fig. 3B,C). The expression level of *Pou5f1* was measured in each cell (Fig.S3B). WT and HET cells assume identity of either epiblast (EPI) or primitive endoderm (PrE): 39 versus 33 respectively (Fig.3A-C Fig.S3A). No MUT cells cluster in proximity of EPI or PrE (Fig.3A, Fig.S3A). ScRNAseq failed to identify any significant expression of maturing PrE markers such as *Sox17, Gata4 or Sox7* (Fig.3D) in MUTs, as predicted from previous IHC or bulk RNA analysis^24, 25^. Rarely, MUT cells expressed *Pdgfrα*, probably reflecting initiation of expression prior to loss of maternal OCT4, since PDGFRα, like GATA6, is a very early presumptive PrE marker^48, 49^.

**Figure 3:**
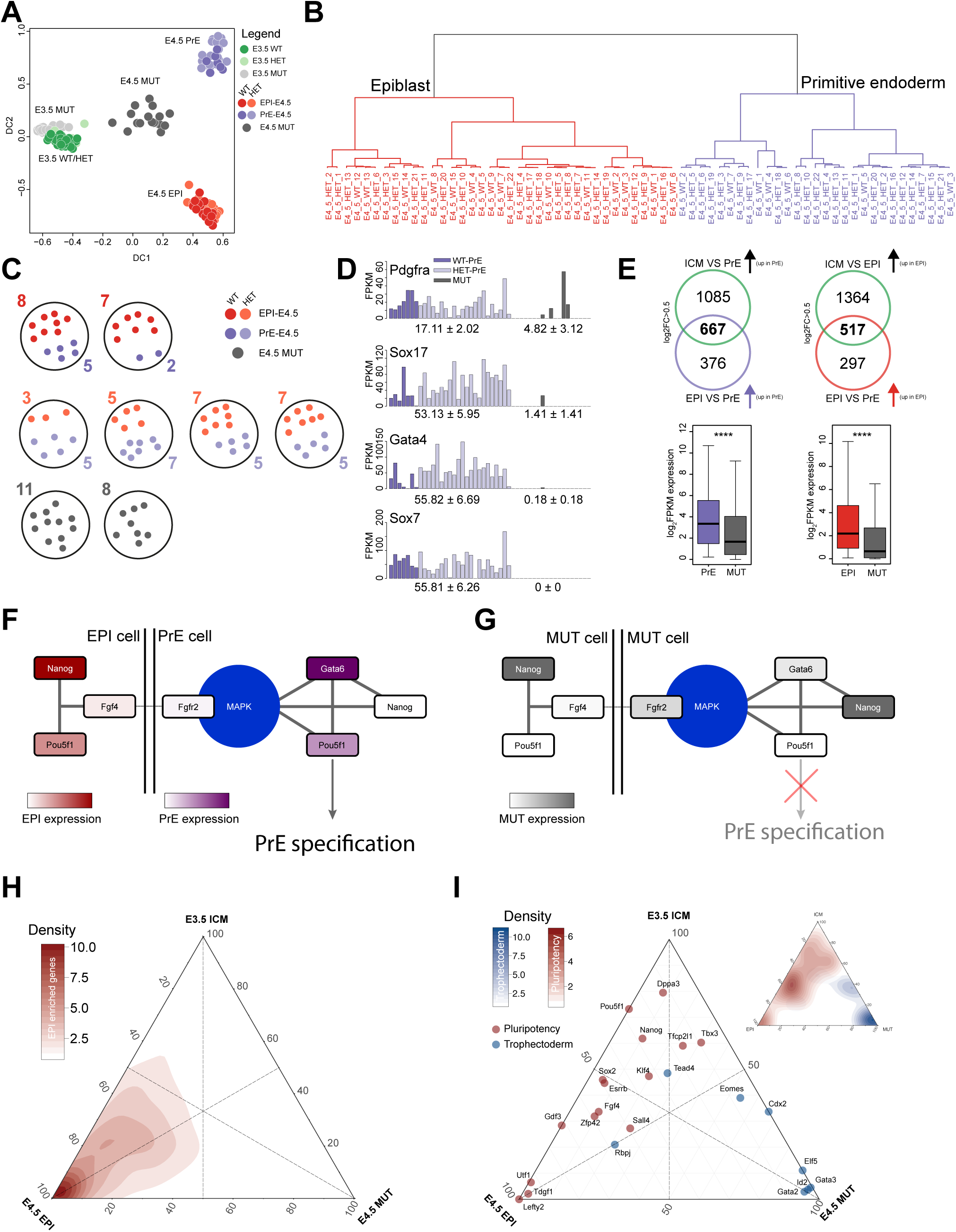
(A) Diffusion plot of early and late blastocyst cells; color represents the different genotypes and lineages. (B) Dendrogram (agglomeration method: ward.D2) for late blastocyst ICM WT/HET cells and (C) schematic representation of number of late ICM single cells per embryo and their genotype. (D) Single cell FPKM expression of PrE markers *(Pdgfra, Sox17, Gata4, Sox7*) in PrE and MUT cells. (E) Top: Venn diagram showing the number of significant (padj<0.05) and enriched PrE and EPI genes. Bottom: Boxplot of log_2_FPKM in late blastocyst PrE and MUT cells of 667 genes and late blastocyst EPI and MUT cells of 517 genes. (F) Network of genes associated with PrE specification in WT cells and (G) mutant cells. (H) Ternary plot of early WT/HET early blastocyst cells, WT/HET EPI and MUT cells. Axes show the relative fraction of expression of 814 EPI enriched genes or (I) pluripotent and trophectoderm associated genes.

WGCNA revealed independent clustering of MUT cells and co-expression of specific genes normally mutually exclusive by E4.5 (Fig.S3C, Suppl.Table4). We assessed quantitatively and qualitatively the PrE and EPI genes underrepresented in MUTs (Fig.3E, Fig.S3D, see methods). A significant drop in intensity in MUT cells was observed, suggesting global failure to activate both PrE and EPI transcription networks. In normal late blastocysts *Gata6* becomes restricted to a subset of cells constituting the PrE. As expected, in WT/HET embryos its expression is mutually exclusive with *Nanog* (Fig.S3E)^49, 50^. However, in MUTs, 7/19 cells co-expressed *Gata6* and *Nanog* (Fig.S3E), suggesting a role for OCT4 in mediating mutual repression, as previously suggested^24^. PrE induction and differentiation is mediated by FGF4 produced from EPI cells^51^ interacting with FGF receptors (R)1 and 2^52, 53^. The failure of this early lineage segregation in MUT ICMs confirms the requirement for OCT4 induction of FGF4^13^ and places this obligation at an early stage in PrE/EPI specification. Consistent with previous findings^13^, E4.5 MUT cells do not express FGF4 but upregulate FGFR1 and FGFR2 (Fig.S3F). We adapted a model of the gene network involved in the second lineage decision^54^ in the WT/HETs compared with MUT cells. In the presence of OCT4, EPI cells express NANOG and FGF4 (Fig.3F). FGF4 drives PrE fate transition and restriction^55^ by triggering ERK signalling, suppressing NANOG and activating PrE markers SOX17, GATA4 and SOX7. However, in MUT cells ERK signal is disrupted and generally downregulated (Fig.S3G) and consequently PrE markers are absent (Fig.3G).

Having identified normal expression of some pluripotency factors in early MUT blastocysts we examined the role of OCT4 in late blastocysts by inspecting expression of EPI-enriched genes (n=814, Fig.3E) and pluripotency markers. Ternary plots represent expression density between three different conditions. We reasoned that if MUT cells fail to express EPI-enriched genes globally, a bias in the density distribution would be expected. Indeed, the EPI/ICM sides of the triangle showed the highest density for EPI enriched genes when compared with MUT (Fig.3H). We then explored how pluripotency and TE associated factors were distributed along the ternary plot. Genes not expressed in MUT cells are located on the axis connecting ICM and EPI; these include *Utf1, Pou5f1, Lefty2* and *Tdgf1*. Overall, most pluripotency factors cluster at the ICM/EPI side, indicating lower but detectable expression in the E4.5 MUT cells (Fig.3I) or TE cells (Fig.S3H). Conversely, genes associated with TE identity, *Gata2, Gata3, Eomes, Id2, Elf5* and the Notch signalling pathway^35, 56–60^ are located on the side specific for MUT (Fig.3I) and TE cells (Fig.S3H). Interestingly we found that *Tead4*, which is a crucial transcriptional regulator of mitochondrial function in TE, is not upregulated in MUT cells, suggesting impairment of mitochondrial function not directly associated with transition of MUT cells to TE (Fig.3I).

### OCT4 MUT cells acquire TE-like identity but exhibit specific differences compared with normal TE

To understand how OCT4 represses TE transcription factors during normal ICM development, we sought to identify specific and common gene expression between WT TE and MUT cells. We consulted published TE single cell data from E3.5 and E4.0 embryos^61^ Diffusion component analysis, coupled with pseudotime reconstruction and non-linear regression, allowed us to identify different developmental trajectories (Fig.4A). Loss of OCT4 and subsequent activation of TE genes drives E4.5 MUT cells towards WT TE. Deconvolution of heterogeneous populations^62^ is an analysis designed to estimate percentage identity of distinctive cells towards a specific endpoint. To quantify similarities between TE and WT/HET/MUT late blastocyst cells we performed quadratic programming to resolve a non-negative least-squares constraint problem (Fig.4B). Similarity between TE and MUT cells was highest, with a median value of ∼0.6 (60%), compared to ∼0.2 (20%) and ∼0.25 (25%) with EPI and PrE cells respectively. We further validated this result with Gene Set Enrichment Analysis (GSEA) by comparing the rank of differentially expressed genes between E4.5 EPI (PrE)/E4.0 TE and E4.5 EPI (PrE)/E4.5 MUT (Fig.S4A,B). Together, these results indicate that late blastocyst MUT cells share a significant portion of the TE transcriptional program. Since our embryos were recovered from nascent implantation sites, they are likely more advanced than those exhibiting non-TE identity profiled previously^24^. We performed a two-way hierarchical analysis with published TE-enriched genes^63^ (Fig.4C,D). Transcripts enriched in early and late TE cells, such as *Krt18, Krt8, Gata3 and Id2*^34, 59, 64, 65^ were also upregulated in MUT cells. Interestingly we also detected co-expression of *Fabp3* and *Cldn4* (Fig.4E). *Fabp3* regulates fatty acid transport in trophoblast cells and plays a central role in fetal development^66^. *Cldn4* is essential for tight junction formation between TE cells during blastocyst formation^67^. We also identified genes expressed in early and late TE not consistently detected in MUT cells, indicating aberrant establishment of the TE transcriptional network. As suggested by pseudotime and diffusion component analysis (Fig.4A), E4.5 MUT cells fail to express a proportion of late TE markers.

**Figure 4:**
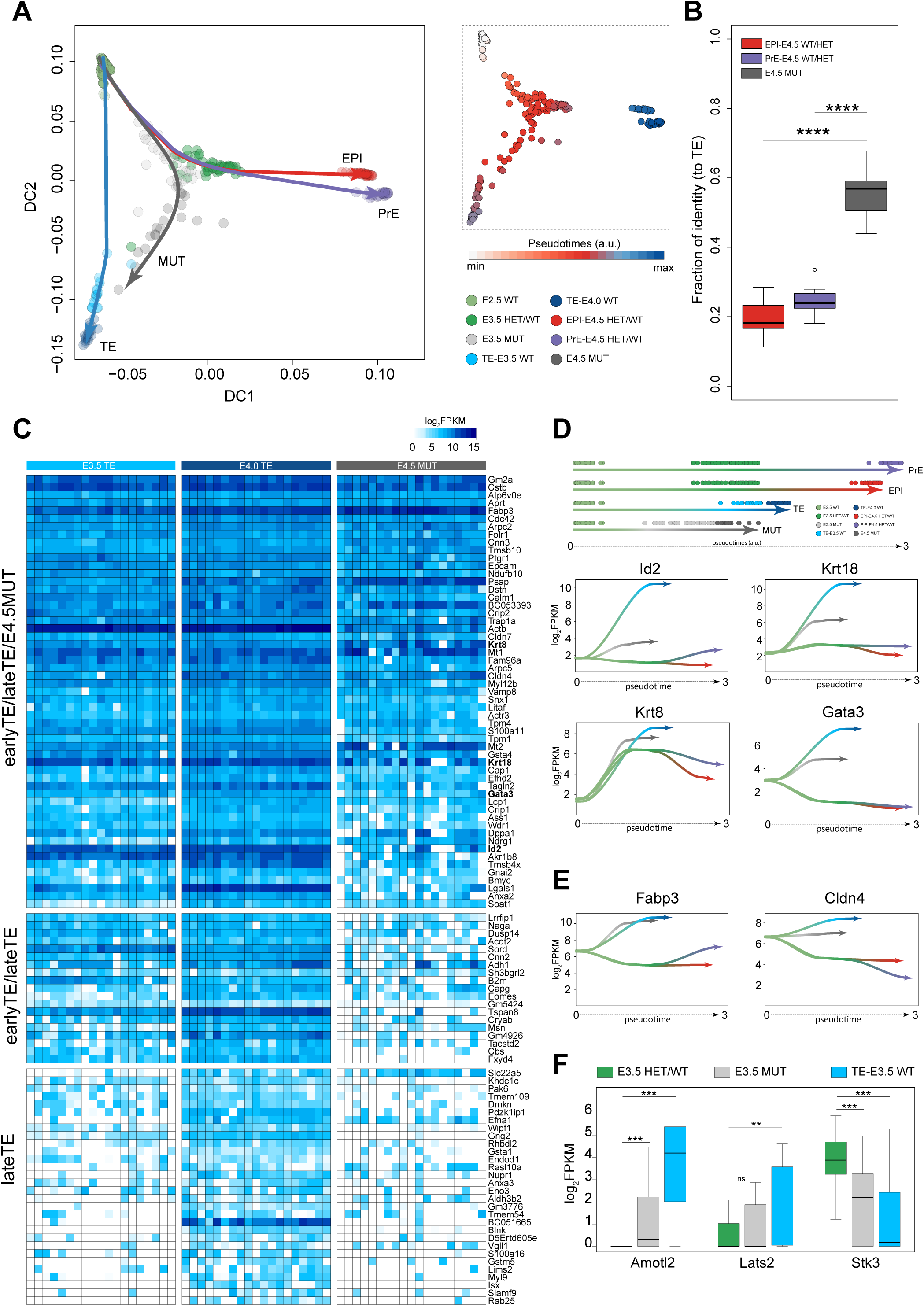
(A) Left panel: diffusion component plot for E3.5/E4.5 WT/HET/MUT cells (this study) and E3.5 and E4.0 TE cells from Deng et al., 2014. Color represents the different genotypes/lineages. Trajectory lines were fitted with cubic line (lambda =0.01). Right panel: diffusion component and pseudotime expression. (B) Fraction of similarities between E4.5 EPI (WT/HET)/E4.5 PrE (WT/HET)/E4.5 MUT and E4.0 TE cells computed using all expressed genes (log_2_FPKM > 0). (C) Heatmap of TE markers identified by Blakeley et al., 2015 between ICM and TE single cells. (D) Identification of lineage trajectories and loess curve fitting between pseudotimes and log_2_FPKM for *Id2, Krt18, Krt8, Gata3*. (E) Loess curve fitting between pseudotimes and log_2_FPKM for *Fabp3* and *Cldn4*. (F) Boxplot of FPKM expression of genes in HIPPO signalling pathway.

HIPPO signalling promotes the first lineage decision in mouse embryos^10, 68^. The three major components of this pathway are the upstream modulators, the core kinase components and the downstream mediators^69^. *Amotl2*, expressed in human and mouse TE and required for TE cell morphology^70^, is also upregulated in MUT cells. Moreover, STK3, a component of the HIPPO core kinase, phosphorylates LATS1/2, which in turn phosphorylates YAP1. Phosphorylated YAP1 is inhibited from entering the nucleus, thus activating HIPPO. Consistent with the role of STK3, AMOTL2 and LATS2 in modulation of the HIPPO pathway, their transcripts were co-expressed in TE and MUT cells (Fig.4F). This suggests a potential role for OCT4 in regulating the balance of HIPPO signalling in MUT cells and therefore initiating the differentiation to TE. MUT cells thus differentiate by expressing a combination of specific early TE transcription factors, signalling pathways and metabolic genes.

### Role of OCT4 in regulation of metabolism

It was previously suggested that OCT4 null embryos exhibit defective metabolism by the mid-late blastocyst stage^24^ and that changes in acetyl-CoA, mediated by glycolysis, control early differentiation^71^. We performed principal component analysis with glycolytic genes. Dimension 1, which explains the largest variability, segregates MUT from EPI/PrE cells (Fig. 5A). The majority of enzymes were downregulated in MUT cells (Fig.5B,C, Fig.S5A) and, interestingly, the rate limiting glycolytic enzymes *Hk2* and *Pkm* together with *Eno1* and *Pgk1* are also putative targets of OCT4 (Fig.5D). KATS enzymes rely on acetyl-CoA, a product of glycolysis, to acetylate the lysine residues on histone proteins and maintain the open chromatin structure associated with pluripotency. We observed a significant downregulation of several KATS enzymes (Fig.5E) in MUT cells, suggesting that OCT4, by regulating glycolysis^72^ indirectly provides sufficient acetyl-CoA to support an open chromatin state.

**Figure 5:**
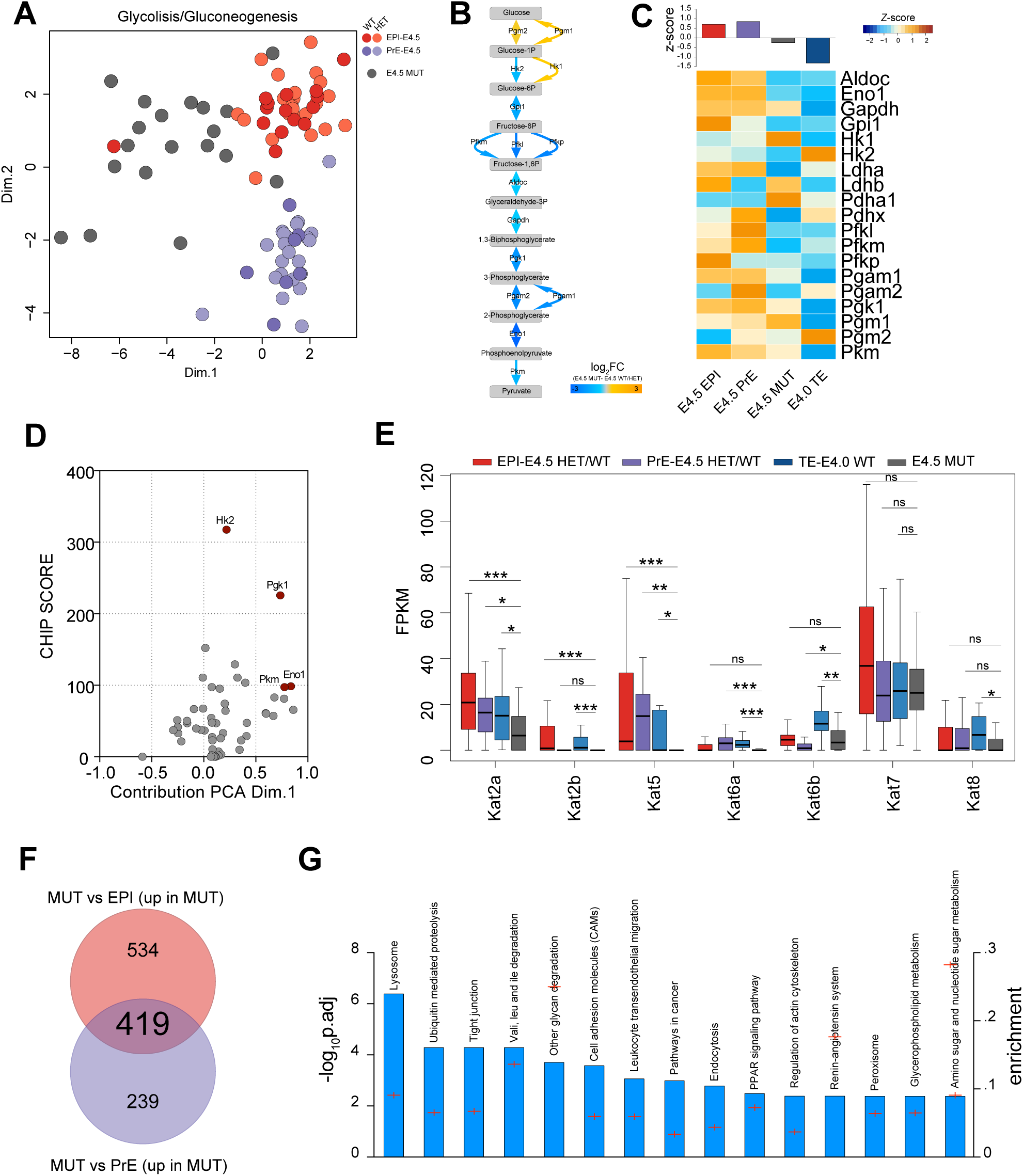
(A) PCA plot of E4.5 WT/HET and MUT cells computed with genes in glycolysis/gluconeogenesis KEGG pathway. (B) Glycolysis pathway with the associated enzymes (arrows) colored by the ratio between E4.5 WT/HET and MUT cells and (C) heatmap of the associated enzymes. (D) Volcano plot showing the contribution on principal component 1 (Fig.5A) and OCT4 CHIPseq score in mESC. (E) Boxplot of Kats gene expression value in E4.5 EPI/PrE WT/HET, E4.0 TE and E4.5 MUT (Student’s t-test; * p<0.05, ** p<0.01, ***p<0.001). (F) Number of variable genes between E4.5 WT EPI/MUT and E4.5 WT PrE/MUT. (G) Enrichment of KEGG pathways computed with 419 common variable genes between MUT and E4.5 EPI/PrE.

To assess systematically the modulated biological processes and pathways we identified 419 common variable genes between E4.5 MUT/E4.5 EPI and E4.5 MUT/E4.5 PrE (Fig.5F) and computed KEGG pathway enrichment (Fig.5G). “Tight junction“, “cell adhesion molecule” and “regulation of actin cytoskeleton” processes suggest that OCT4 regulates important components of biophysical properties of ICM cells. Interestingly, the most significant enriched process was “Lysosome”, indicating a strong and pivotal role of this pathway in MUT cells. Accordingly, processes related to “Lysosome” were also significantly enriched, including “Peroxisome”, “Glycerophospholipid Metabolism”, “Endocytosis”, “PPAR signalling pathway” and “Vali, leu and ile degradation”.

### TFEB translocates into the nucleus in cells lacking OCT4

To determine whether the activation of the lysosomal pathway was a TE characteristic we explored differentially expressed genes and found that only MUT cells, but not WT TE, upregulated a significant proportion of lysosomal genes (Fig.6A). Lysosome is essential for recycling, recruitment of lipids via autophagy and hydrolases, for redistribution of catabolites to maintain cellular function^73^. Autophagy is a catabolic response to starvation^74^. Most autophagy-related genes, such as *Atg,* were upregulated in MUT cells (Fig.6B). Moreover, MUT cells undergo a significant upregulation of fatty acid degradation genes (Fig.6C). The master regulator of lysosomal biogenesis and autophagy is TFEB^74^. TFEB is dissociated by inactive mTORC1 and migrates into the nucleus to activate lysosomal/autophagy genes. The positive regulator of mTORC1 (*Rptor*) is downregulated in MUT cells and, consistently, we found upregulation of *Deptor*, a known negative regulator of mTORC1^75^ (Suppl.Table5). To confirm activation of the lysosomal pathway via TFEB, we performed IHC on *Oct4* conditionally deleted ESC. In OCT4-positive cells, TFEB is localized mainly in the cytoplasm. After OCT4 deletion, a significant translocation of TFEB from cytosol to nucleus occurs (Fig.6D, Fig.S6A). Together, these results indicate that, in response to an altered and energy insufficient metabolism, MUT cells upregulate lysosomal and autophagy pathways to provide cellular energy.

**Figure 6:**
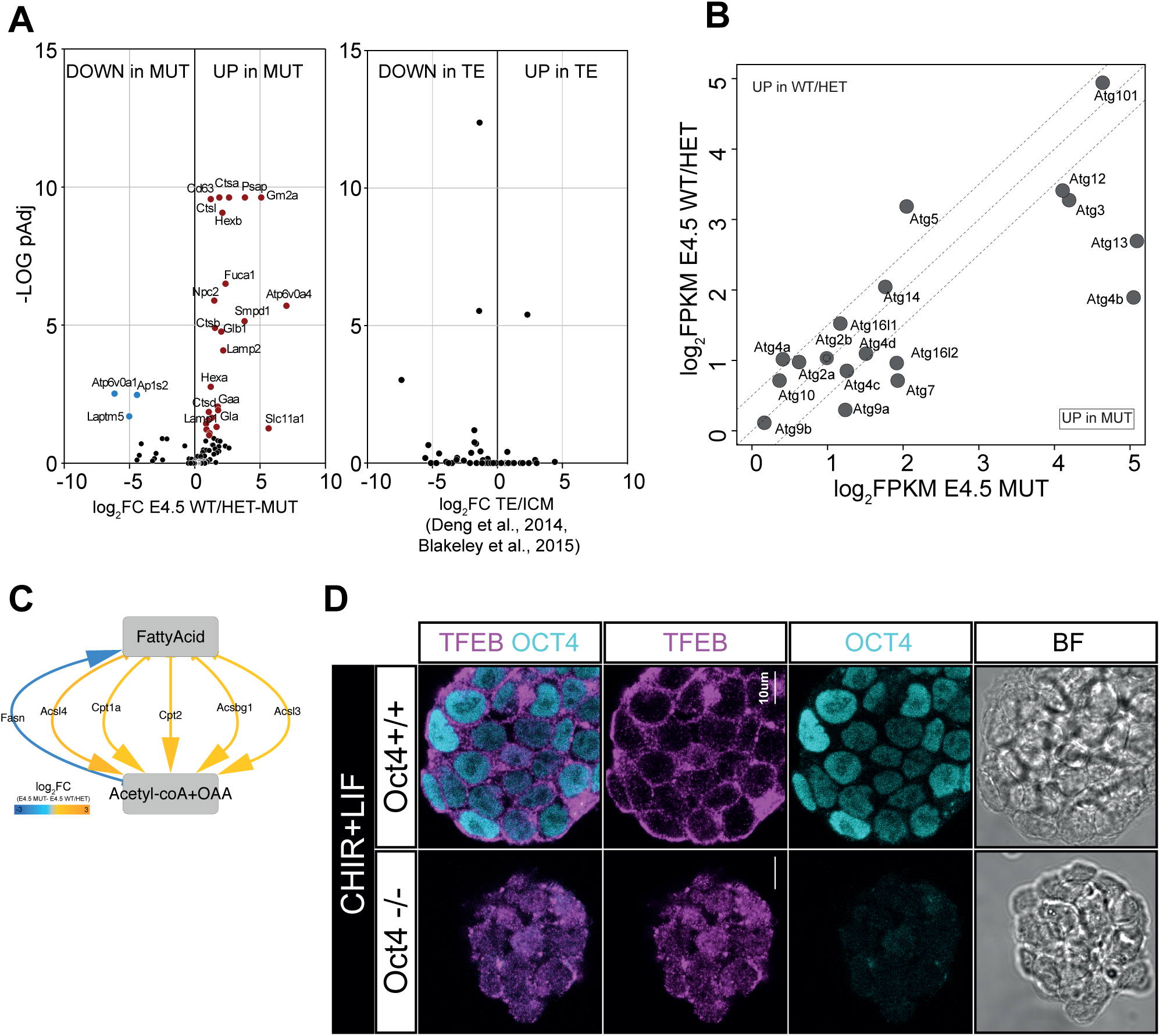
(A) Volcano plot of lysosomal genes variable between E4.5 WT/HET and E4.5 MUT and between WT TE and WT ICM (Blakeley et al., 2015). (B) Scatter plot of autophagy related genes (*Atg*s) between E4.5 WT/HET ICM and E4.5 MUT. (C) Enzymes in fatty acid oxidation and synthetase; color represents the ratio between E4.5 WT/HET and MUT cells. (D) TFEB localization in OCT4+/+ and OCT4-/- cells cultured in CHIR+LIF.

## Discussion

Apart from the known direct targets of OCT4, such as *Utf1*^40^ expression of most other pluripotency-associated factors, including the essential embryonic pluripotency factors NANOG, SOX2 and ESRRB, is not significantly different between MUT cells and WT/HETs at the mid blastocyst stage (E3.5) at both the mRNA and protein level. Detection of most pluripotency-associated factors in OCT4 MUT mid blastocysts suggests independence from OCT4 at this stage, confirming that the state of naïve pluripotency, as captured in the form of ESCs *in vitro*, is not yet attained by the ICM, as reported previously^76^.

Down-regulation of *Utf1* suggests an indirect role for OCT4 in governing epigenetic structure of pluripotent cells, which may account for the precocious expression of some TE factors in MUT cells, preceding changes in expression of most pluripotency factors. Another putative OCT4 target, *Il6st*, is a co-receptor essential for STAT3 signalling in ESCs^77^. We observed significant downregulation of STAT3 target genes in MUT cells as well as reduced P-STAT3 protein and its pluripotency-associated targets, TFCP2L1 and KLF4^45, 46^. Diversion of ICM cells to TE has been observed in a proportion of embryos following maternal/zygotic deletion of *Stat3,* which was attributed to loss of activation of *Oct4*^43^. The diminution of *Socs3* exhibited by MUT cells is an expected direct consequence of reduced STAT3 activity.

Signalling pathways related to matrix organization, including regulation of actin cytoskeleton and cell adhesion molecules are significantly affected in MUT cells. Such processes are associated with exit from pluripotency^78^; cytoskeletal conformational changes inducing cell spreading are associated with differentiation. Our results therefore implicate OCT4 as a mediator for regulation of the biophysical properties of undifferentiated cells.

In this study we dissected the role of metabolism in OCT4 MUT cells. We linked the reduction of glycolysis with the downregulation of most *Kats* enzymes. KATS enzymes rely on acetyl-CoA, a product of glycolysis, to acetylate the lysine residues on histone proteins and maintain an open chromatin structure, associated with pluripotency. *Kats* enzymes are lower in MUT cells, suggesting that OCT4, by regulating glycolysis^72^, indirectly provides sufficient acetyl-CoA to support an open chromatin state. We revealed that most enzymes in glycolytic pathways are downregulated in MUT cells. This may be because some of the rate-limiting enzymes (*Hk2, Pgk1, Pkm* and *Eno1)* are putative targets of OCT4. We also noted downregulation in MUT cells of genes associated with cell respiration. This is likely to be a downstream effect of reduced STAT3 signalling in MUT cells, consistent with the recent report that STAT3 promotes oxidative respiration for maintenance and induction of pluripotency^79^ (Fig. 7A,B). Consequently, respiration processes are disrupted in OCT4 MUT cells where STAT3 is strongly downregulated.

**Figure 7:**
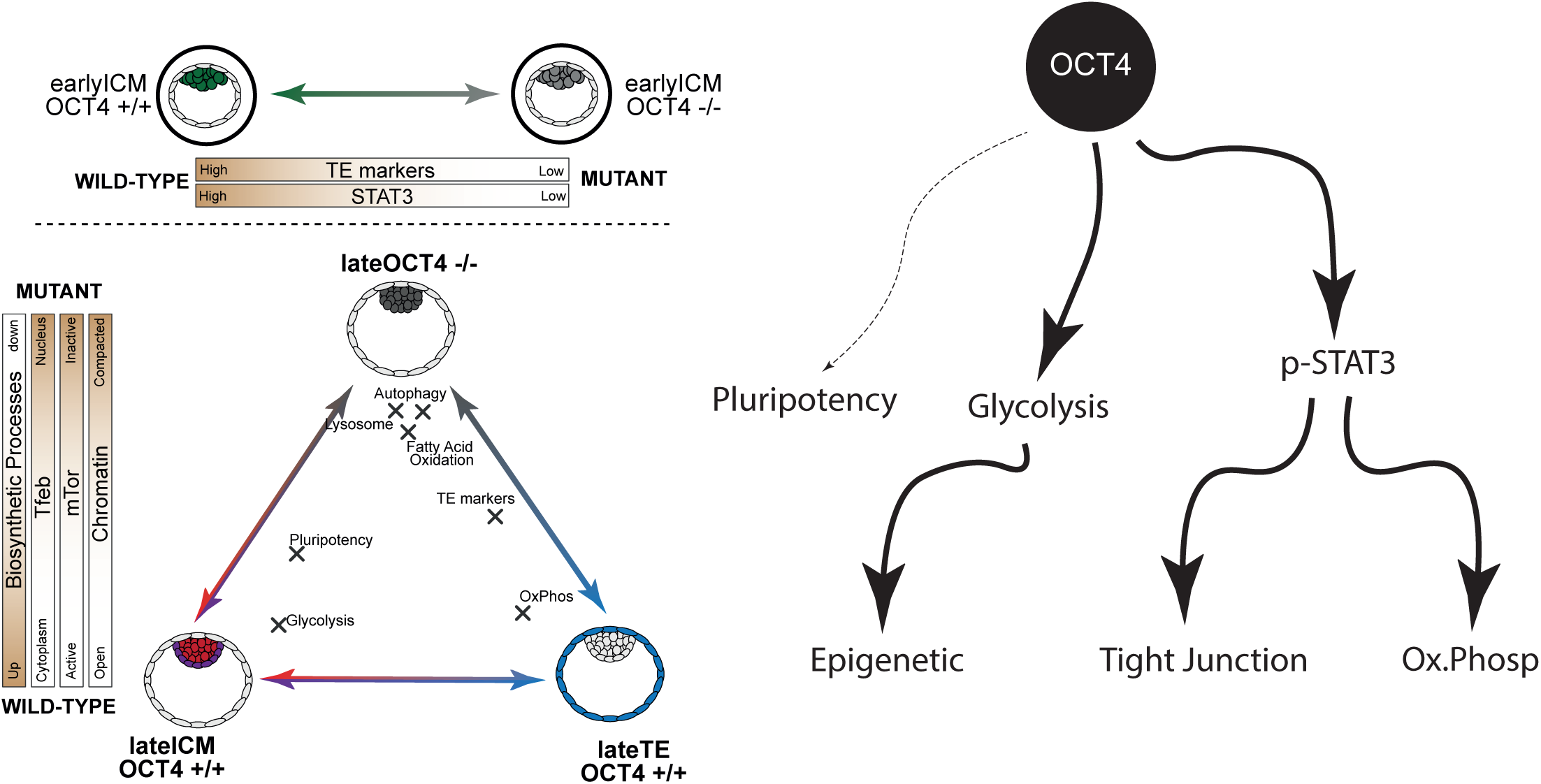
Scheme of OCT4 function in pre-implantation embryo development.

Our scRNAseq data indicates that TFEB pathway is specifically activated in MUT cells as they transition to TE. We provide functional evidence for the requirement for OCT4 in lysosomal activity by demonstrating TFEB translocation into the nucleus during conditional deletion of *Oct4* in ESCs. We propose that MUT cells upregulate lysosomal gene expression and autophagy to counteract the downregulation of glycolysis and the tricarboxylic acid cycle. Thus, OCT4 is required to maintain the metabolic state needed for survival of pluripotent cells.

In summary, our systematic analysis at the single cell level reveals an *in vivo* function for OCT4 in regulating metabolic and biophysical cellular properties via energy metabolism, cell morphology and chromatin accessibility for establishment of pluripotency in the developing mouse embryo (Fig.7).

## Acknowledgements

We are grateful to Lawrence Bates, Kevin Chalut, Rosalind Drummond, Peter Humphreys, Kenneth Jones, Masaki Kinoshita, Carla Mulas and Maike Paramor for material, intellectual and technical contribution to the project. This work was supported by the University of Cambridge, BBSRC project grant RG74277, MRC PhD studentships for AK and HS and a core support grant from the Wellcome Trust and MRC to the Wellcome Trust – Medical Research Council Cambridge Stem Cell Institute.

## Author contributions

JN, AK and GGS conceived the project; AK, AY, HS, TEB and JN carried out experiments; GGS performed data analysis; AK and SES performed imaging analysis; all authors contributed to writing the manuscript.

## Competing interests

The authors have no competing interests

## Supplementary figures

**Figure S1:**
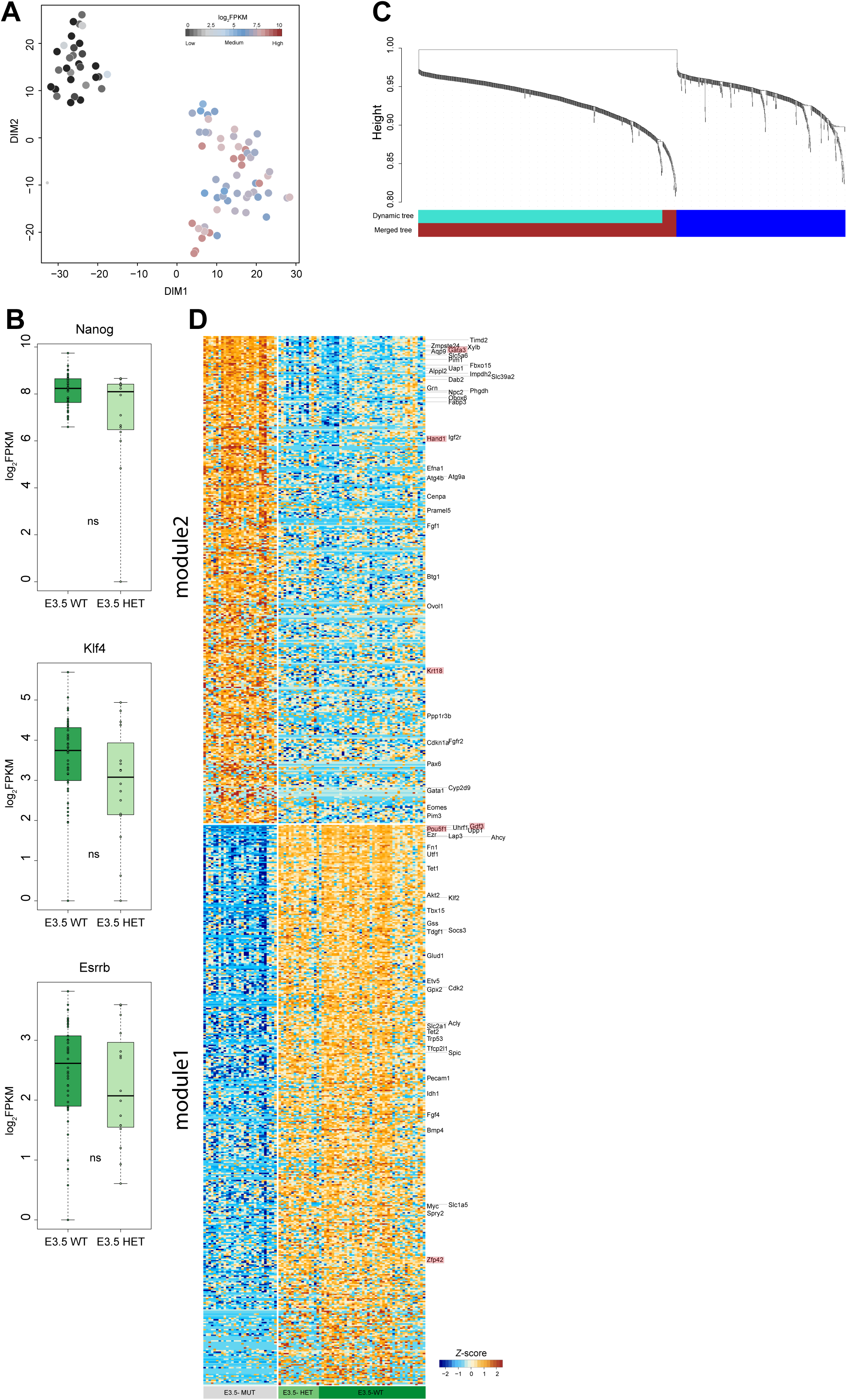
(A) t-SNE plot for early blastocyst cells. Sample colors represent *Pou5f1* log_2_ expression. (B) Barplots for *Nanog*, *Klf4* and *Esrrb* expression. (C) Genes dendrogram computed with WGCNA. (D) Heatmap showing the top co-regulated genes between E3.5 MUT and E3.5 WT/HET (> 5 interaction; n=346 MUT genes and n=398 WT/HET genes).

**Figure S2:**
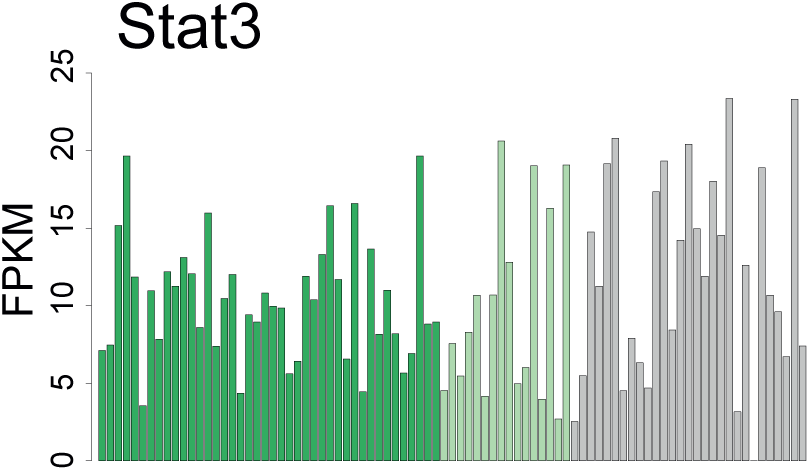
Barplot of total *Stat3* expression for E3.5 WT, HET and MUT cells.

**Figure S3:**
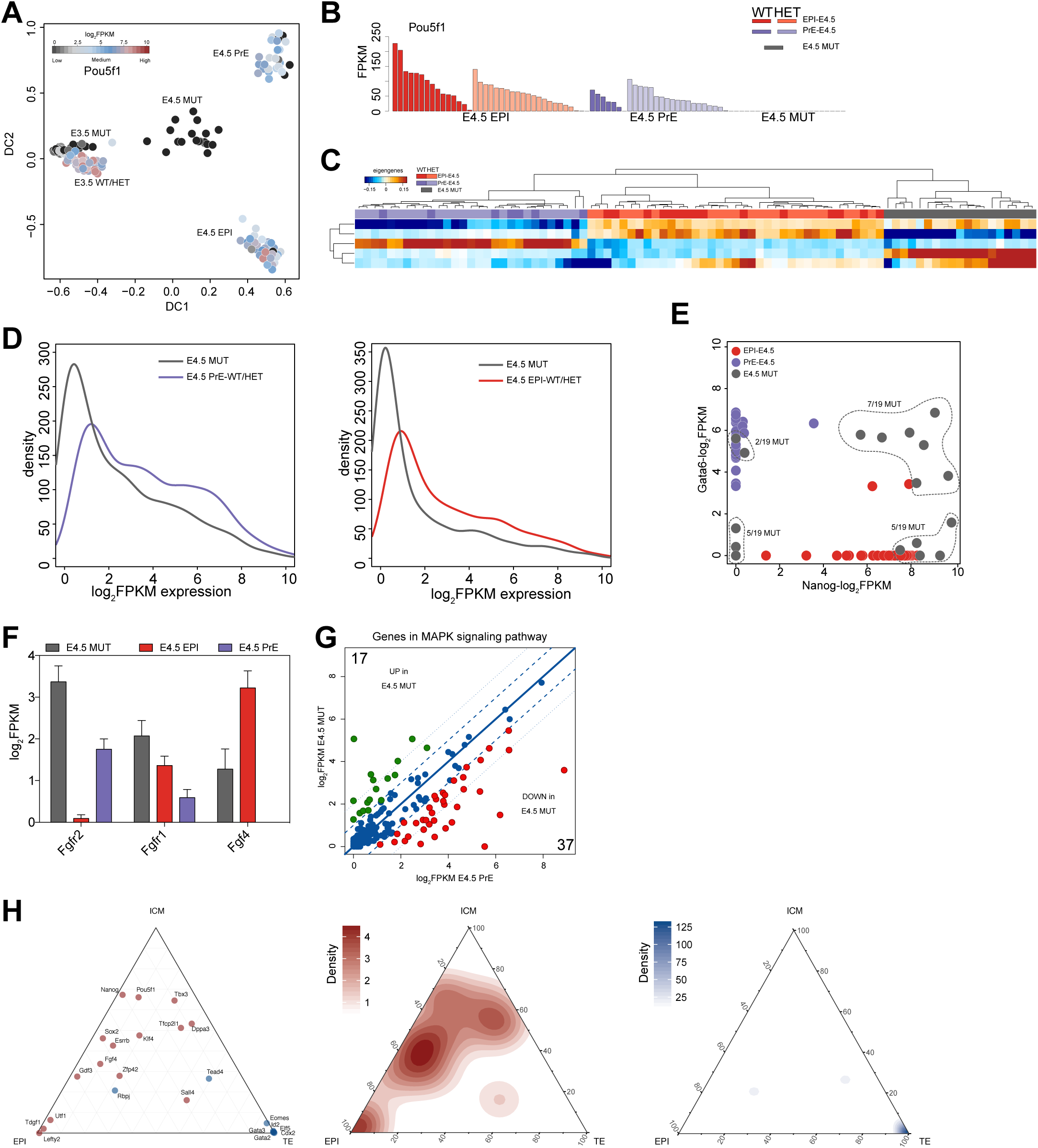
(A) Diffusion plot of early and late blastocyst cells; color represents *Pou5f1* expression. (B) Barplot shows FPKM expression of *Pou5f1* for each single cell at E4.5 stage. (C) One-way hierarchical cluster of eigengenes value computed from WGCNA (power 8; dist=0.35, size =30) (D) Density distribution of log_2_FPKM in late blastocyst PrE and MUT cells of 667 genes and late blastocyst EPI and MUT cells of 517 genes. (E) Scatter plot of *Nanog* and *Gata6* FPKM expression values for E4.5 EPI (WT/HET), E4.5 PrE (WT/HET) and E4.5 MUT cells. (F) log_2_FPKM expression of Fgf4, Fgfr1 and Fgfr2 in WT/HET EPI/PrE and MUT cells. (G) log_2_FPKM scatter plot of MAPK signalling genes between late PrE and MUT blastocyst cells. (H) Ternary plot of early WT/HET blastocyst cells, WT/HET EPI and TE cells. Axes show the density of the relative fraction of expression.

**Figure S4:**
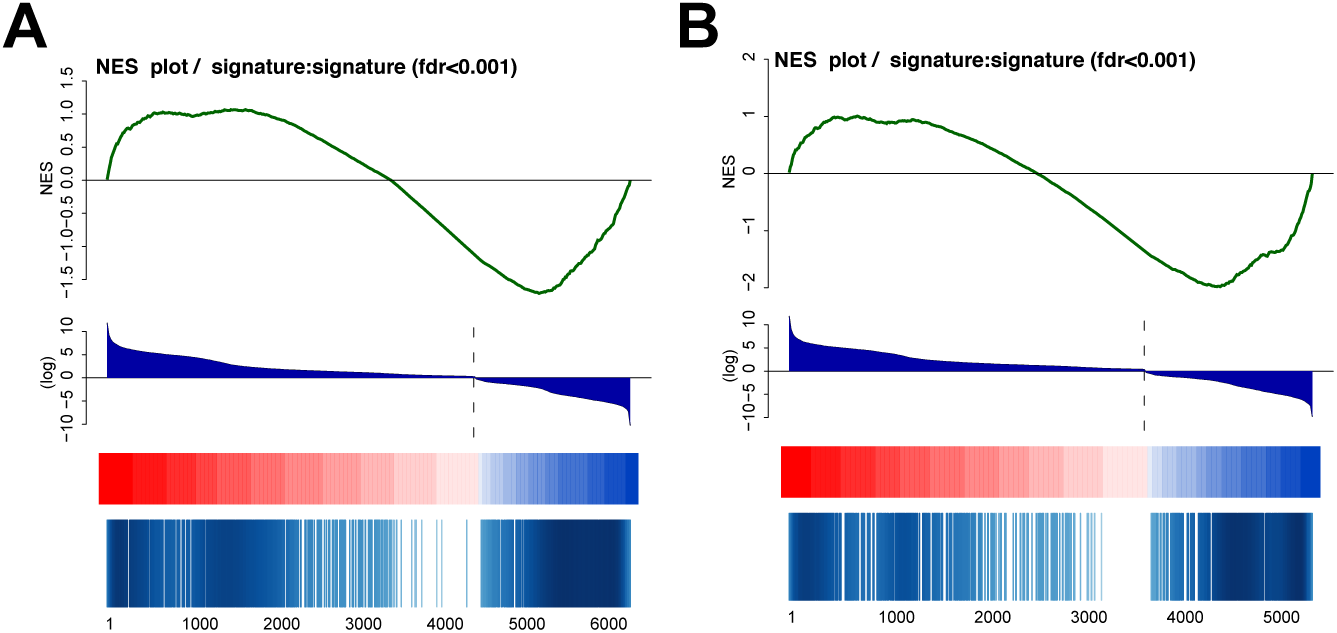
(A) GSEA analysis computed with log_2_FC between E4.5 EPI-WT.HET/E4.0 TE and (B) E4.5 MUT/E4.0 TE.

**Figure S5:**
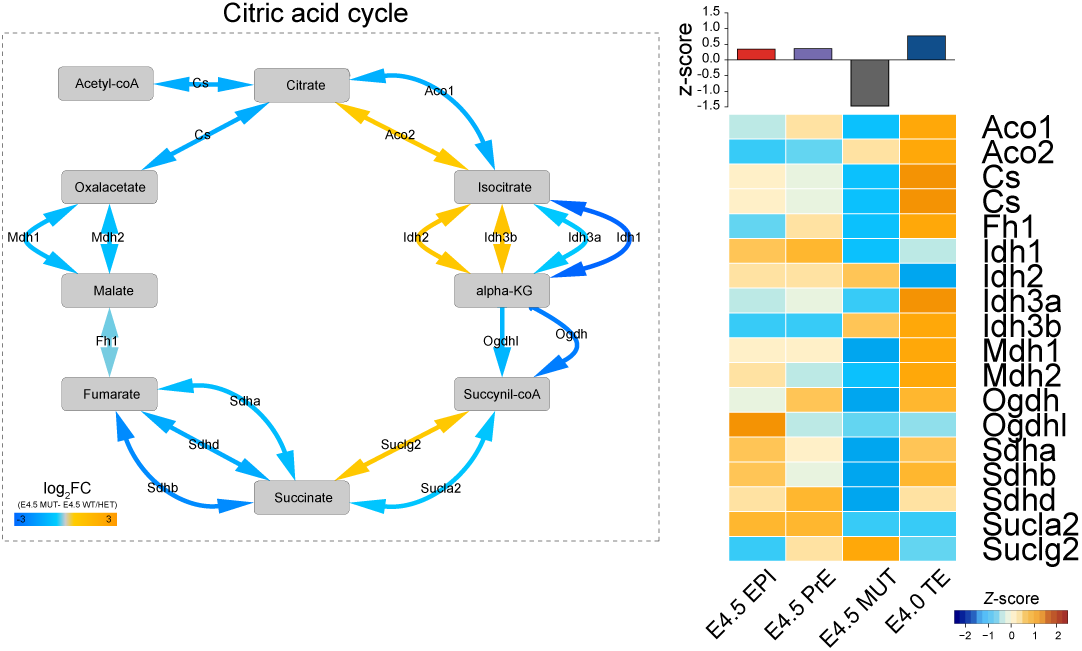
TCA associated enzymes (arrows) color by the ratio between E4.5 WT and MUT cells and heatmap of the associated enzymes.

**Figure S6:**
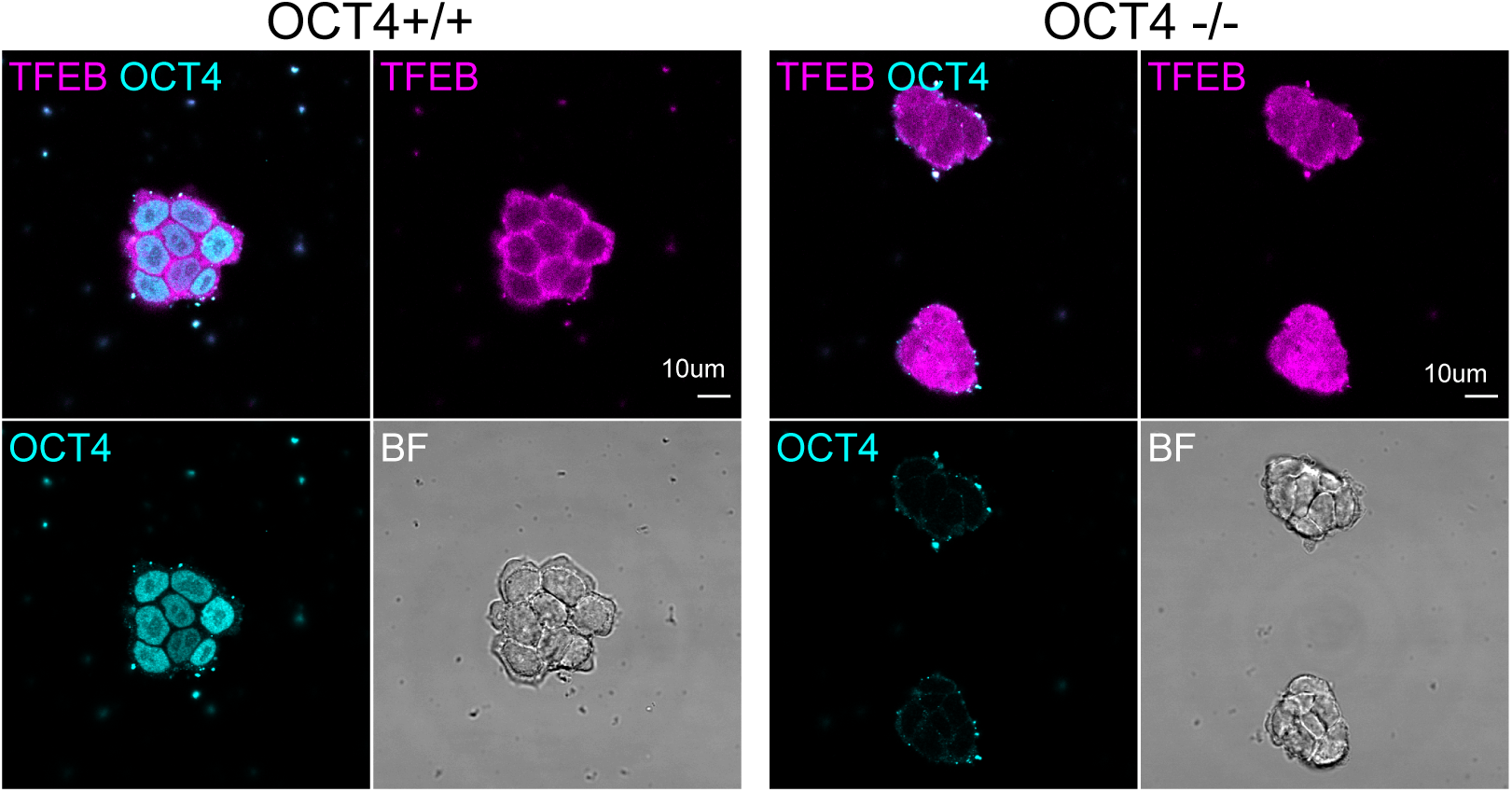
TFEB localization in OCT4+/+ and OCT4-/- cells cultured in CHIR+LIF.

## Materials and Methods

Experiments were performed in accordance with EU guidelines for the care and use of laboratory animals and under the authority of appropriate UK governmental legislation. Use of animals in this project was approved by the Animal Welfare and Ethical Review Body for the University of Cambridge and relevant Home Office licences are in place.

### Mice and husbandry

All embryos were generated from transgenic mouse strains with mixed genetic backgrounds. They were: Oct4_+/-1_, ZP3CreTg_/+2_, R26::CreERT2_3_ and Oct4_LoxP/LoxP4_. Compound transgenic mice were generated from crosses of these lines. Genotyping was performed by PCR analysis using DNA extracted from ear biopsies or trophectoderm lysate following isolation of ICMs by immunosurgery^1, 5^. Primer sequences are as follows:

Oct4LoxP: CTCAAACCCCAGGTGATCTTCAAAAC; GGATCCCATGCCCTCTTCTGGT
Oct4 null: GCCTTCCTCTATAGGTTGGGCTCCAACC; GGGCTGACCGCTTCCTCGTGCTTTACG; GAGCTTATGATCTGATGTCCATCTCTGTGC
Cre transgene: GCGGTCTGGCAGTAAAAACTATC; GTGAAACAGCATTGCTGTCACTT

Amplification was carried out on around 5 µL of lysate for 35 cycles (following 95°C hot start for 10 minutes) of 94°C, 15 seconds; 60°C, 12 seconds; 72°C, 60 seconds, with a final extension at 72°C for 10 minutes. Reaction products were resolved by agarose gel electrophoresis. Mice were maintained on a lighting regime of 14:10 hours light:dark with food and water supplied *ad libitum*. Embryos for RNAseq were generated from Oct4_+/-_ *inter se* natural mating; those for IHC were compound transgenics derived from Oct4_LoxP/-_; ZP3Cre_Tg/+_ stud males and Oct4_LoxP/LoxP_ dams. Detection of a copulation plug following natural mating indicated embryonic day (E) 0.5. Embryos were isolated in M2 medium (Sigma) at E3.5 or E4.5.

### ESCs and culture

Indole-3-acetic acid (IAA, Sigma) inducible Oct4 deletable induced pluripotent stem cells (iPSC) were kindly provided by Lawrence Bates. Neural stem cells (NSCs) derived from brains of E13.5 Oct4_fl/-_ Rosa26::CreER embryos, as previously reported^6^ were treated with 500 nM 4-hydroxytamoxifen to induce deletion of the floxed *Oct4* allele. Oct4-/- NSCs were transduced with retroviral Oct4, Klf4 and cMyc. Retroviruses were produced in PLAT-E cells; briefly, cells were transfected with pMXs-Oct3/4, pMXs-Klf4 and pMXs-cMyc using FuGENE 6 reagent (Promega). Polybrene (Sigma) was added to a final concentration of 4 μg/ml. 24 hours later NSCs were nucleofected with pPB-CAG-Oct4AID-PGK-hygro (Piggybac transposon containing a constitutive Oct4AID fusion protein expression cassette and a constitutive hygromycin resistance expression cassette), pPB-CAG-Tir1-IRES-bsd (Piggybac transposon containing a constitutive Tir1 IRES blasticidin resistance expression cassette) and pPBase (a non-integrating Piggbac transposase expression vector) using Amaxa Nucleofection Technology (Lonza AG) according to the manufacturer’s instructions. Program T-020 was used for NSC nucleofections. Cells were plated in NSC medium for 2 days then switched to medium (GMEM [Sigma] containing 10% FCS [Sigma], 1× NEAA [PAA], 1× penicillin/streptomycin [PAA], 1 mM sodium pyruvate [PAA], 0.1 mM 2-mercaptoethanol [Gibco] and 2 mM L-glutamine [Gibco], supplemented with 20 ng/ml LIF). Medium was switched to KSR medium (GMEM containing 10% KSR [Invitrogen], 1% FCS, 1× NEAA, 1× penicillin/streptomycin, 1 mM sodium pyruvate, 0.1 mM 2-mercaptoethanol, 2 mM L-glutamine) supplemented with 2iL (20 ng/ml LIF, 3 μM CHIR99021 [CHIR], and 1 μM PD0325901 [PD03]), and selection was added for expression of the endogenous (floxed) Oct4 locus on the ninth day in KSR-2iL. Expanded colonies were passaged into N2B27 + 2iL. 0.8 µg of pPB-CAG-GFP-IRES Zeocin (gift from Masaki Kinoshita) and 0.4 µg of pPy-CAG Pbase were transfected into IAA inducible Oct4 deletable iPSCs using lipofectamine 2000 (Thermo Fisher Scientific). The transfected cells were picked after selection with Zeocin (100mg/ml) and expanded. The resulting iPSCs were routinely maintained on 0.1% gelatin (Sigma)-coated 6-well plates (Falcon) in N2B27 + 2iL and passaged every three days following dissociation with Accutase.

### Cell differentiation

IAA inducibly deletable Oct4 cells were seeded (1.5 x 10^4^) on fibronectin-coated (12.5µg/ml; Millipore) Ibidi-dishes (µ-Dish, 35mm) and cultured in N2B27 + 2iL for one day. The next day, medium was switched to N2B27 + 100U/ml LIF, 3 µM CHIR and 500 µM IAA for Oct4 deletion (or 0.1% ethanol for controls) and cells were cultured for another day.

### Immunohistochemistry

#### Embryos

The zona pellucida was removed from all non-hatched embryos using acid tyrodes solution (Sigma). Embryos were fixed in 4% PFA (paraformaldehyde; Thermo Fisher Scientific) in PBS at room temperature for 15 minutes. After rinsing in PBS/PVP (3mg/ml PVP in PBS) they were permeabilised in 0.25% Triton X (Sigma) in PBS/PVP for 30 minutes, then incubated in 2% donkey serum, 0.1% BSA and 0.01% Tween 20 in PBS (blocking buffer) for at least 15 minutes and incubated overnight at 4°C in primary antibodies diluted in blocking buffer (Suppl. Table 7). After 3 x 15 min washes in blocking buffer they were incubated at room temperature in the dark in secondary antibody (Alexa dye, Life Sciences) and DAPI (4’,6-diamidino-2-phenylindole/ Invitrogen) 1:500 in blocking solution. After 3 x 15 minute washes, the embryos were taken through 25%, 50%, 75% then 100% VectaShield mounting medium (Vector Laboratories) in blocking solution. The embryos were placed in a drop of Vectashield on coverslips and surrounded by spots of Vaseline on which the upper coverslip was gently pressed. To avoid drying out the coverslip was surrounded by nail polish. Images were acquired using a Leica TCS SP5 confocal microscope.

#### Cells

Oct4-deleted and control ESCs were fixed with 4% PFA in PBS at room temperature for 15 minutes, then rinsed in PBS and blocked in PBS containing 3% donkey serum (Sigma), 0.1%TritonX at 4°C for 2-3 hours. Primary antibodies (Suppl. Table 7) were diluted in blocking buffer, and samples were incubated in the appropriate antibody solution at 4°C overnight. They were rinsed three times in PBST, compromising PBS + 0.1% TritonX, for 15 minutes each. Secondary antibodies were diluted in blocking buffer with or without 500 ng/ml DAPI and samples were incubated in the appropriate secondary antibody solution at room temperature for 1 hour in the dark. They were rinsed three times in PBST for 15 minutes each, then stored in PBS at 4°C in the dark until imaging.

#### Imaging

Samples were observed using a spinning disk microscope (Andor Revolution XD System with a Nikon Eclipse Ti Spinning Disk) or a Leica TCS Sp5 confocal. 40x objective lens was used with Type F immersion liquid. The images were analysed by Fiji as described previously^7^.

#### Preparation of samples for RNA-sequencing

For E3.5 blastocysts, zona pellucidae were removed using acid tyrode’s solution (Sigma) and embryos subjected to immunosurgery^1, 5^ using 20% anti-mouse whole antiserum (Sigma) in N2B27 in at 37°C, 7% CO_2_ for 30 minutes, followed by 3 rinses in M2, then 15 minutes in 20% non-heat inactivated rat serum (made in house) in N2B27 in at 37°C, 7% CO_2_. After 30 minutes in fresh N2B27 lysed trophectoderm was removed and placed in lysis buffer for genotyping. ICMs were incubated in 0.025% trypsin (Invitrogen) plus 1% chick serum (Sigma) for 5-10 minutes in small drops and dissociated by repetitive pipetting using a small diameter mouth-controlled flame-pulled Pasteur pipette. Individual ICM cells were transferred into single cell lysis buffer and snap frozen on dry ice. Smart-seq2 libraries were prepared as described previously^8^ and sequenced on the Illumina platform in a 125 bp paired-end format.

#### RNA-seq data processing

Early/mid and late blastocyst annotated cell data was downloaded from GSE45719 and selected only cells expressing trophectoderm markers^9^. Genome build GRCm38/mm10 and STAR 2.5.2a^10^ were used for aligning reads and ensembl release 87^11^ was used to guide gene annotation. After removal of inadequate samples according to filtering criteria described^12^, we quantified alignments to gene loci with htseq-count^13^ based on annotation from Ensembl 87.

#### Transcriptome analysis

Principal component and cluster analyses were performed based on log_2_FPKM values computed with custom scripts, in addition to the Bioconductor packages *DESeq*^14^ or *FactoMineR*. Diffusion maps and T-distributed stochastic neighbor embedding (t-SNE) were produced with *destiny*^15^ and *Rtsne* packages.

Default parameters were used unless otherwise indicated. Differential expression analysis was performed with *Single Cell Differential Expression R package, scde*^16^, which has the advantage of fitting individual error models for the assessment of differential expression between sample groups. For global analyses, we considered only genes with FPKM > 0 in at least one condition, not expressed genes were always omitted. Euclidean distance and average agglomeration methods were used for cluster analyses unless otherwise indicated. Expression data are made available in Supplemental Tables and through a web application to visualise transcription expression and fitted curve with temporal pseudotime of individual genes in embryonic lineages (https://giulianostirparo.shinyapps.io/pou5f1/). High variable genes across cells were computed according to the methods described^12, 17^. A non-linear regression curve was fitted between average log_2_ FPKM and the square of coefficient of variation (log CV_2_); then, specific thresholds were applied along the x-axis (average log_2_ FPKM) and *y*-axis (log CV_2_) to identify the most variable genes.

To assess the accuracy of the identified lineages, we used the Weighted Gene Co-Expression Network Analysis unsupervised clustering method (WGCNA^18^ to identify specific modules of co-expressed genes in each developmental lineage/genotype. R package ggtern was used to compute and visualize ternary plots. Kyoto Encyclopedia of Genes and Genomes (KEGG) was used to compute pathway enrichment and to download genes in glycolysis/gluconeogenesis and tricarboxylic acid cycle pathways.

#### Quadratic programming

Fractional identity between pre-implantation stages was computed using R package DeconRNASeq^19^. This package uses quadratic programming computation to estimate the proportion of distinctive types of tissue. The average expression of pre-implantation stages (E4.5 WT/HET epiblast and primitive endoderm, E4.5 MUT cells) were used as “signature” dataset.

Finally, the fraction of identity between TE cells and the “signature” dataset was computed using the overlapping gene expression data (FPKM > 0).

## Data availability

GEO submission

## Code availability

Code is available upon request

